# Chromes from Chromatin: Sonification of the Epigenome

**DOI:** 10.1101/037523

**Authors:** Davide Cittaro, Dejan Lazarevic, Paolo Provero

## Abstract

The epigenetic modifications are organized in patterns determining the functional properties of the underlying genome. Such patterns, typically measured by ChIP-seq assays of histone modifications, can be combined and translated into musical scores, summarizing multiple signals into a single waveform. As music is recognized as a universal way to convey meaningful information (1), we wanted to investigate properties of music obtained by sonification of ChIP-seq data. We show that the music produced by such quantitative signals is perceived by human listener as more pleasant than that produced from randomized signals. Moreover, the waveform can be analyzed to predict phenotypic properties, such as differential gene expression.

**Significance Statement:** Music is recognized as universal way to communicate emotions and, more in general, meaningful information. Various sources of information can be translated into music or sounds, mostly for recreational purposes. It has been shown that human ear can classify information encoded into sounds. Quantitative genomic features, and in particular epigenetic marks, do represent functional information that is exploited by cells to drive biological processes. We test a method to translate such information into music and we study some properties of the sonificated chromatin marks. We show that not only musical representation of epigenetic marks has intrinsic musicality, but also that differences in musical representation of genomic loci reflect differences of the RNA levels of the underlying genes.

## Introduction

Sonification is the process of converting data into sound. Sonification itself has a long, yet punctuated, story of applications in molecular biology, several algorithms to translate DNA (2) or protein sequences (3, 4) to musical scores have been proposed. The same principles have also been extended to the analysis of complex data (5) showing that, all in all, sonification can be used to describe and classify data. Indeed, the very same procedures may also be applied for recreational purposes.

One of the limitations of sonification of actual DNA and protein sequences is their intrinsic conservative nature. Assuming the differences in two individual genomes are, on average, one nucleotide every kilobase (6), the corresponding musical scores would have little differences.

On the contrary, dynamic ranges typical of transcriptomic and epigenomic data may provide a richer source for sonification.

In this work we describe an approach to convert ChIP-seq signals, and in principle any quantitative genomic feature, into a musical score. We started working on our approach for amusement mainly, and we realized that the sonificated chromatin signal were surprisingly harmonious. We then tried to assess some properties of the music tracks we were able to generate. We show that the emerging sounds are not random and instead appear more melodious and tuneful than music generated from randomized notes. We also show that different ChIP-seq signals can be combined into a single musical track and that tracks representing different conditions can be compared allowing for the prediction of differentially expressed genes.

Examples of sonification for various genomic loci are available at https://soundcloud.com/davide-cittaro/sets/k562

## Definitions

MIDI: MIDI (Musical Instrument Digital Interface) is a standard that describes protocols for data exchange among a variety of digital musical instruments, computers and related devices. MIDI format encodes information about note notation, pitch, velocity and other parameters controlling note execution (*e.g.* volume and signals for synchronization).

MIDI file format: a binary format representing MIDI data in a hierarchical set of objects. At the top of hierarchy there is a Pattern, which contains a list of Tracks. A Track is a list of MIDI events, encoding for note properties. MIDI events happen at specific time, which is always relative to the start of the track. MIDI Resolution: resolution sets the number of times the status byte is sent for a quarter note. The higher the resolution, the more natural the sound is perceived. Resolution is the number of Ticks per quarter note. At a specific resolution *R,* Tick duration in microseconds *T*is related to tempo (expressed in Beats per Minute, BPM) by the following equation

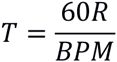

## Results

### Approach

In order to translate a single ChIP-seq signal track to music we bin the signal over a specified genomic interval *(i.e.* chrom:start-end) into fixed-size windows (*e.g.* 300 bp) and note duration will be proportional to the size of such windows. As we are dealing with MIDI standard, we let the user specify track resolution and the number of ticks per window (see definitions); the combination of these parameters defines the duration of a single note. The default parameters associate a bin of 300 bp with one quaver (1/8 note).

In order to define the note pitch, we take the logarithm of the average intensity of the ChIP-seq signal in a genomic bin. The sounding range of the whole signal is discretized in a predefined number of semitones. At default parameters, the range is binned into 52 semitones, covering four octaves. In order to introduce pauses, the lowest bin of the signal range represents a rest. If two consecutive notes or rests fall in the same bin, we merge them in one note doubling its duration.

Using this approach, any ChIP-seq signal can be mapped to a chromatic scale. We implemented the possibility to map a signal on a different scale (major, minor, pentatonic…); to this end, intensity bin boundaries are merged according to the definition of a specific scale (Figure 1). MIDI tracks produced in this way can be then imported into a sequencer software where they can be further processed, setting tempo and time signature.

**Figure 1:**
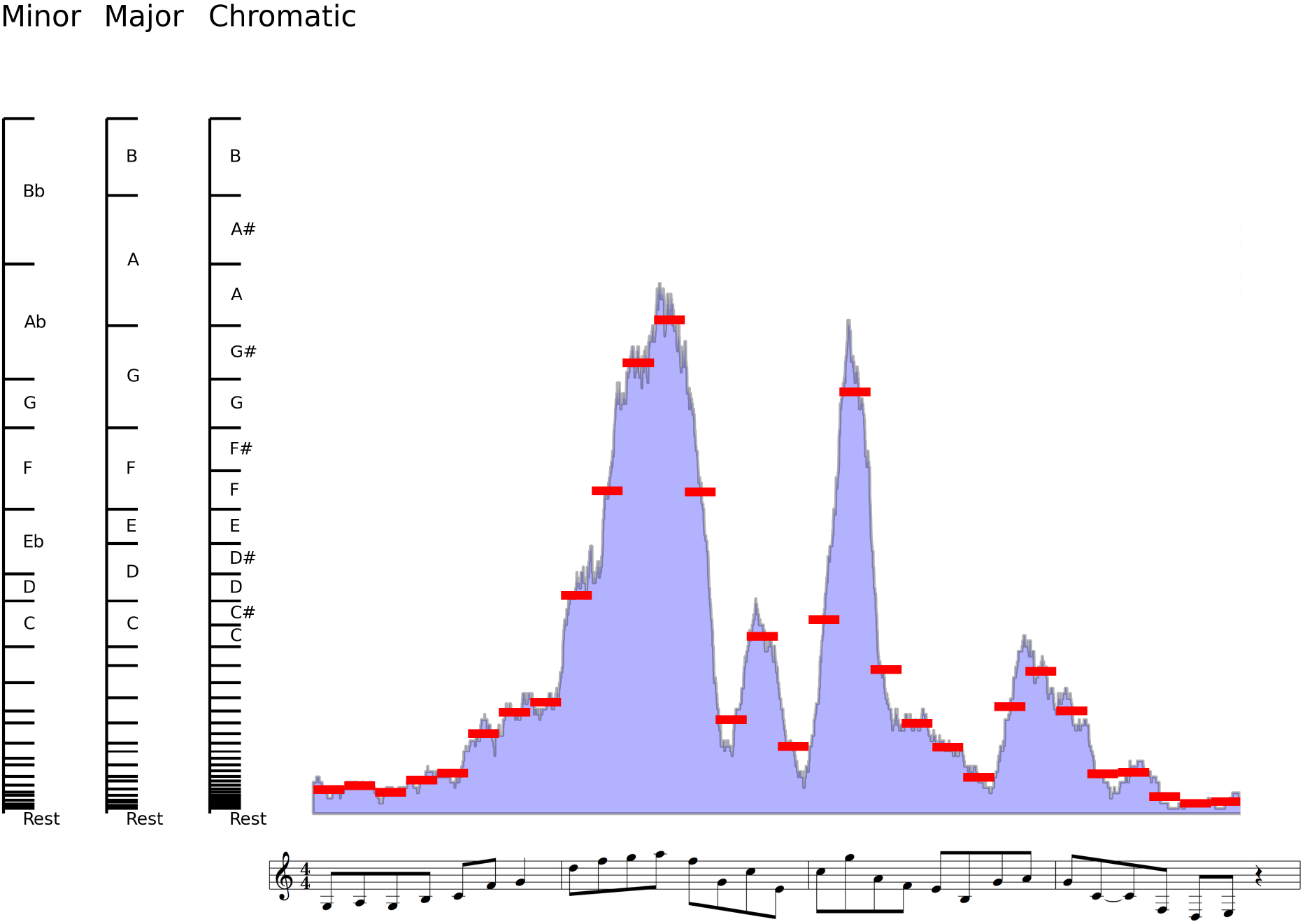
Graphical representation o the approach used to transform quantitative signals to music. ChIP-seq values (H3K4me3 in the example) are binned in fixed-size intervals over the genome. Each interval corresponds to a 1/8 note. Average values of log-transform of read counts in each genomic bin (red lines) are matched into predefined number of semitones (chromatic scale). Notes may be mapped to a specified scale (major and minor scales are exemplified in the figure). Consecutive equal notes are merged in single note with double duration. Values falling in the first bin are considered rests.

### Music produced from chromatin marks is not perceived as a random pattern

In order to test whether sonification of chromatin marks are perceived as random patterns, we selected ten genomic regions and generated corresponding tracks based on the following histone modifications: H3K27me3, H3K27ac, H3K9ac, H3K36me3, H3K4me1, H3K4me2, H3K4me3, H3K9me3 (Supporting Audio files S1.1 to S10.1). For the same regions, we randomized genomic signal at base and bin level (Supporting Audio files S1.2 to S10.2). When data are randomized at base level, the average intensity is uniforms across the bins, resulting in a repeated note (data not shown); this is largely expected as ChIP-seq signals are distributed on the genome according to a Poisson law (7) or, more precisely, to a Negative Binomial law (8).

Randomization at bin level, instead, equals to shuffling notes during the execution. We administrated a questionnaire to a set of volunteers *(n=8)* not previously tested for education in music. Volunteers were asked to listen to each pair of original/random track and choose which track they felt was more appealing. Track order was randomized when testing different volunteers. Notably, in the majority of the cases (62/80) the music generated from genomic signal without randomization was judged more appealing. Results are significant to a Fisher-exact test (*p*=1.95e-3), suggesting that genomic signals contain information that can be recognized by human ear. The number of correct answers for each volunteer ranged from 5 to 10, with a median value of 8.

### Differences in musical tracks reflect differences in gene expression

Once we assessed the existence of musical pattern in genomics signals, we were keen to explore if this kind of information could be exploited to identify biological features of samples. Since the epigenetic DNA modifications reflected by histone marks influence gene expression (9), we tested if differences in musical tracks generated from various ChIP-seq signals reflects differences in gene expression of the corresponding loci. To this end, we downloaded ChIP-seq marks (H3K27me3, H3K27ac, H3K9ac, H3K36me3, H3K4me1, H3K4me2, H3K4me3, H3K9me3, Pol2b) and RNA-seq data for K562 and NHEK cell lines from the ENCODE project (10). For each RefSeq locus we converted ChIP-seq signals to music with fixed parameters (see methods). RNA-seq data were used to identify genes that are differentially expressed between the two cell lines, under a *p*-value < 0.01 and | logFC|>1, according to recent SEQC recommendations (11).

A common way to classify music is based on summarization of track features after spectral analysis (12, 13). Such approach involves the summarization of track as Mel-Frequency Cepstral Coefficients (MFCC) that are subsequently clustered using Gaussian Mixture Models (GMM). A distance between tracks can then be defined as described in (14), who used it as a classifier for musical genres.

We tested if a similar approach could be used to develop a predictor of differential expression based on the distance between musical tracks generated from two cell lines.

We defined a distance between songs as described in methods and we optimized the parameters using as a training set the 250 genes with the most significant differential expression *p*-value and as many genes with the least significant *p*-value according to RNA-seq (Figure 2). We found that optimal performance is at MFCC=30 and GMM=10, with an AUC=0.609.

**Figure 2:**
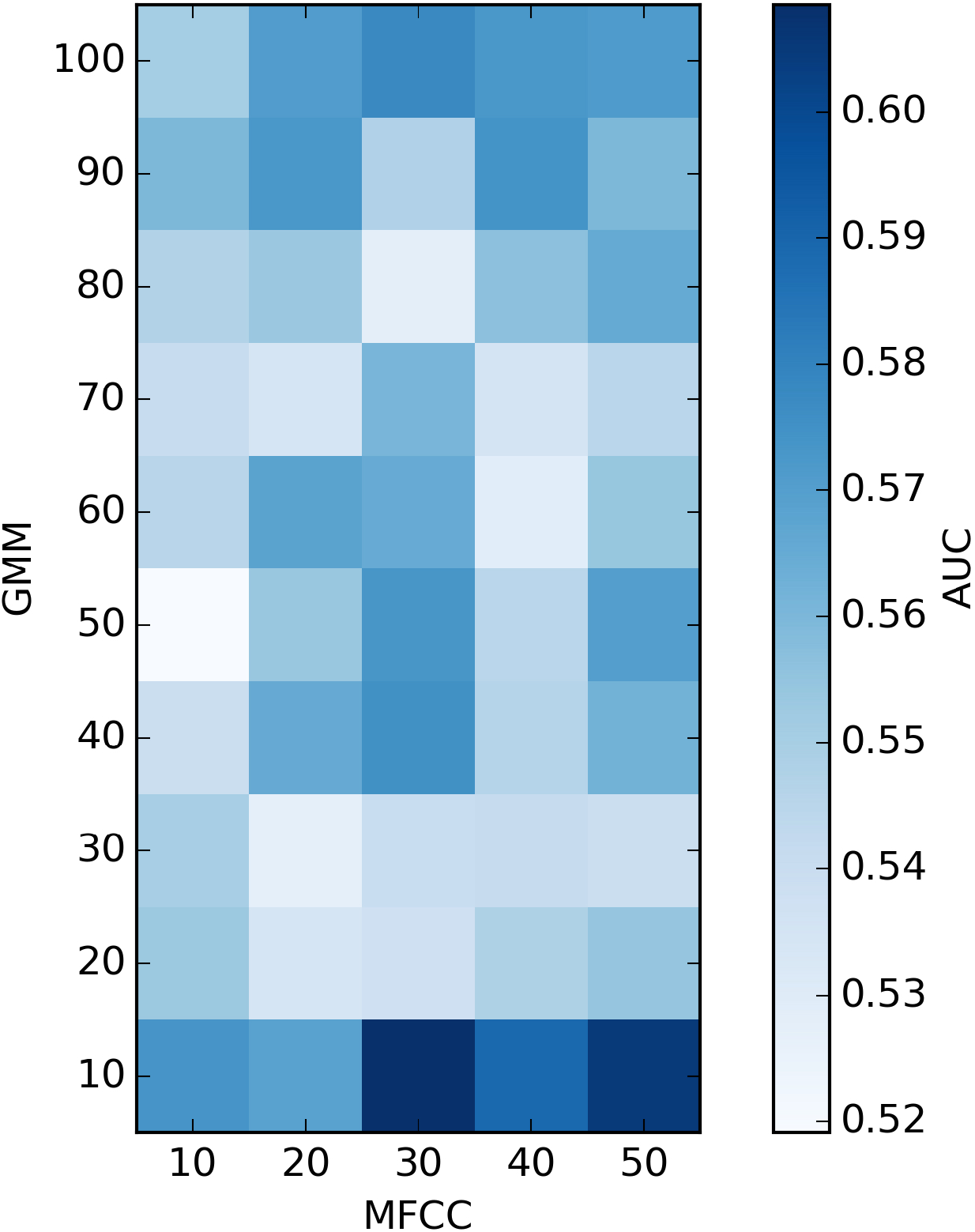
Evaluation of different combination of parameters in predicting differential gene expression on a gold-standard subset of 500 genes. Each square corresponds to a number of Mel-Frequency Cepstral Coefficients (MFCC) used to summarize signal and a number of centers for Gaussian Mixture Model (GMM). Colors are given by the corresponding Area Under the Curve (AUC)

We summarized tracks representing all RefSeq genes using such parameters, we then compared distances with differential expression performing a ROC analysis. Our results indicate that differences in information contained in musical representation of chromatin signals may be linked to differential expression, although power of prediction is limited (AUC=0.5184, *p*=1.4597e-03).

### Similarity between musical tracks overlaps similar biological properties

As additional issue we wanted to assess if similarities between musical representation of chromatin status may be linked to the biology of the underlying genes. To this end, we calculated pairwise distances for all regions using parameters identified above on K562 cell line. Hierarchical clustering of the distance matrix identifies eight major clusters (Figure 3, left). We performed Gene Ontology Enrichment analysis on each cluster, here represented as word cloud of significant terms (Figure 3, center, supplementary Table S2); we found that different clusters are linked to genes showing different biological properties. For example, some clusters (6, 7 and 8) were linked to regulation of cell cycle, others were linked to metabolic processes (2 and 5) or vesicle transport (3 and 4). We also evaluated the distribution of expression (expressed as log(RPKM)) of the underlying genes (Figure 3, right); we found that regions clustered by the distance between musical tracks broadly reflects groups of genes with different level of expression, spotting clusters of higher expression (cluster 5) or lower expression (clusers 2 and 3); assessment of statistical significance of differences in distribution of gene expression values among clusters is presented in Table 1.

**Table 1.** p-values of Mann-Whitney U-test for differences in distribution of expression between clusters. Tests showing significant difference (p ≤ 0.05) are presented in boldface.

|  | *Cluster 2* | *Cluster 3* | *Cluster 4* | *Cluster 5* | *Cluster 6* | *Cluster 7* | *Cluster 8* |
| --- | --- | --- | --- | --- | --- | --- | --- |
| *Cluster 1* | **2.635e-23** | **3.548e-18** | **8.127e-37** | **8.627e-125** | **2.859e-83** | **3.231e-71** | **3.169e-63** |
| *Cluster 2* |  | 1.068e-01 | 2.008e-01 | 4.005e-01 | 2.801e-01 | 7.907e-02 | 9.467e-02 |
| *Cluster 3* |  |  | 2.880e-01 | **2.225e-02** | 1.399e-01 | 3.910e-01 | 3.745e-01 |
| *Cluster 4* |  |  |  | **4.159e-02** | 3.025e-01 | 2.614e-01 | 2.821e-01 |
| *Cluster 5* |  |  |  |  | **5.098e-04** | **3.898e-09** | **7.586e-07** |
| *Cluster 6* |  |  |  |  |  | **1.259e-02** | **2.866e-02** |
| *Cluster 7* |  |  |  |  |  |  | 4.545e-01 |

**Figure 3:**
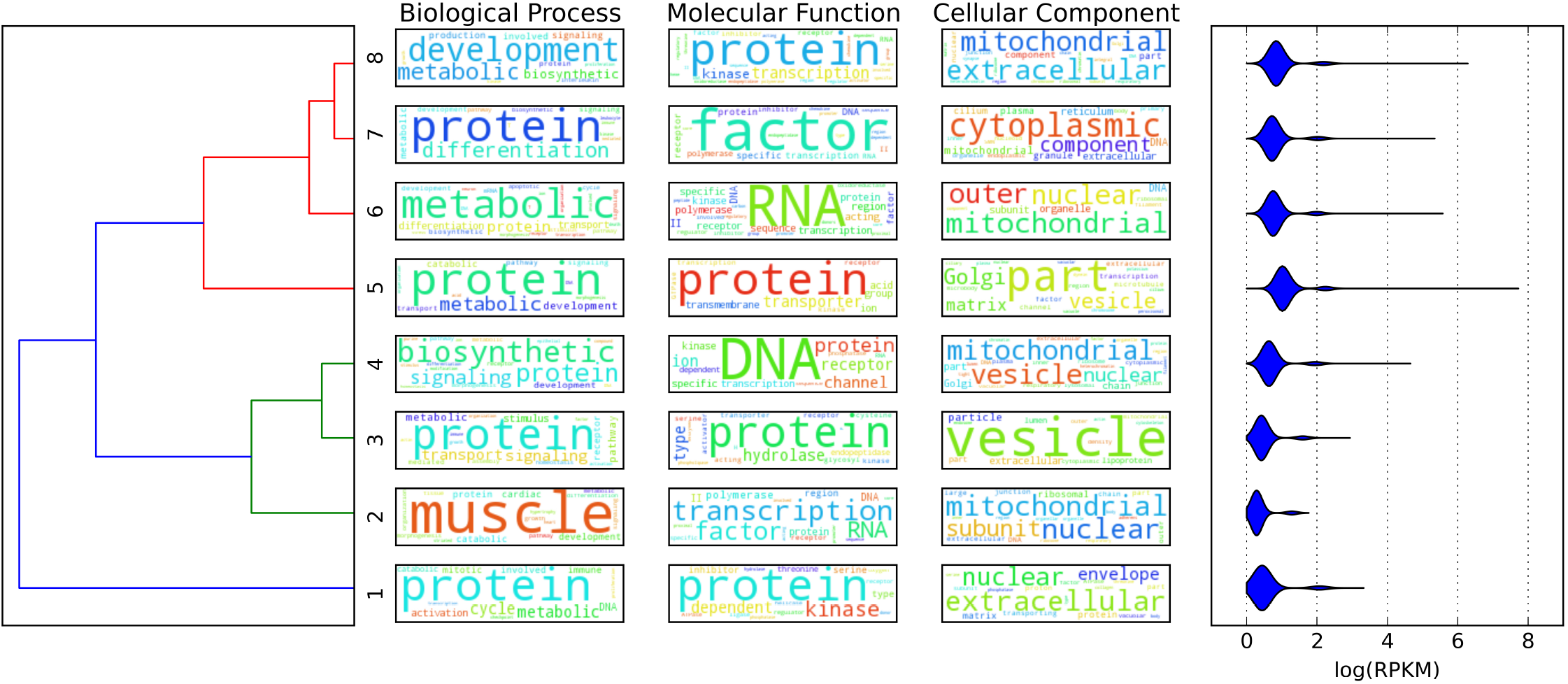
Hierarchical clustering of genomic regions identifies 8 main clusters (left). Each cluster broadly corresponds to specific biological properties according to Gene Ontology enriched terms (middle). Level of expression of genes included in each cluster show specific distributions (right).

## Discussion

Chromatin shape and genome function are governed, among several factors, by the coordinated organization of epigenetic marks (15). Modifications of such marks are dynamic and are fine-tuned during the life of a cell or an organism. Analysis of histone modifications, as well as transcription factors and other proteins binding DNA, by ChIP-seq already described patterns of enrichment that are specific to their relative function (16, 17). Analysis of combinatorial patterns of histone modifications already unveiled its potential in understanding functional properties of the genome (18,19) and the cross-talk among multiple chromatin marks (20).

We show, in this work, that the information carried by multiple histone modifications can be caught in a human-friendly way by translating ChIP-seq signals into musical scores. Although the investigation of the psychological factors that underlie tuneful perception of sonificated genomic signals is out of the scope of this manuscript, our results suggest that human hearing is able to perceive patterns conveying information encoded in ChIP-seq data analyzed and to distinguish from random noise.

We automated the analysis of differences between musical tracks using an established method based on summarization of spectral data. By this approach, we investigated the possible link between differences in ways chromatin sounds and phenotypic features. Our results suggest that differences in transcript levels can be predicted by the differences of sonificated genomic regions, although performances of such approach are limited. We reasoned that many factors may explain such poor results: first of all there is a vast space of parameters that can be tuned to create a single musical track and we still lack methods to explore it efficiently. In addition, the Mel scale used to summarize audio signal has been developed to match human capabilities to perceive sound (21), hence it may not be optimal for the comparison of the tracks generated in this work.

It has already been shown that it is possible to predict levels of gene expression starting from chromatin states, although the method used to perform chromatin segmentation has a large impact on such predictions (22). In this work we found that differences in chromatin-derived music reflects, to some extent, differences in level of expression of underlying genes and their related biology.

To conclude, although we cannot advocate the usage of musical analysis as universal tool to analyze biological data yet, we confirm that quantitative features on the genome are patterned and contain information, hence can be converted into sounds that are perceived as musical. We limited our analysis on specific chromatin modifications, but in principle any quantitative genomic feature can be converted and integrated into a musical track. The choice of parameters and instruments has been standardized for the analysis presented, for illustrative purpose we show that different signals from the same region can be combined using different instruments (https://soundcloud.com/davide-cittaro/random-locus-blues) and signals from different genomic regions can be merged (https://soundcloud.com/davide-cittaro/non-homologous-end-joining).

## Materials and Methods

### Sonification of ENCODE ChIP-seq data

Raw data for various modifications were downloaded from (GSE26320). Read tags were aligned to human genome (hg19) using bwa aligner (23). Alignments were converted to bigwig tracks (24) after filtering for duplicates and quality score higher than 15.

In order to define regions to be converted to music scores, we selected intervals around RefSeq gene definition, from 1kb upstream of TSS to 2kb downstream of TES. ChIP-seq signal were firstly converted to MIDI using custom scripts (https://bitbucket.org/dawe/enconcert) according to parameters defined in Table 2. MIDI tracks belonging to the same region from the same sample were merged into a single MIDI file, converted to WAV format using timidity software (http://timidity.sourceforge.net), with the exception of tracks presented as supplementary information which have been processed with Apple GarageBand software.

**Table 2.** Parameters used to convert different ChIP-seq signals into corresponding musical trakcs

| **Antibody** | **Scale** | **Octave** | **Key** | **Tick Size** | **Bin Size** |
| --- | --- | --- | --- | --- | --- |
| *H3K27me3* | *minor* | *4* | *B* | *600* | *400* |
| *H3K27ac* | *minor* | *3* | *B* | *900* | *600* |
| *H3K9ac* | *minor* | *3* | *B* | *1200* | *800* |
| *H3K36me3* | *minor* | *4* | *B* | *300* | *200* |
| *H3K4me1* | *minor* | *3* | *B* | *1200* | *800* |
| *H3K4me2* | *minor* | *3* | *B* | *300* | *200* |
| *H3K4me3* | *minor* | *3* | *B* | *300* | *200* |
| *H3K9me3* | *minor* | *4* | *B* | *300* | *200* |
| *Pol2* | *minor* | *4* | *B* | *600* | *400* |

### Comparison of WAV tracks

In order to compare four samples for each converted genomic region, we extracted MFCC using python_speech_features library (https://github.com/jameslyons/python_speech_features). Selected components were then clustered using Gaussian Mixture Models, implemented in scikit-learn python library 0.15.2 (http://scikit-learn.org). Distance between two tracks were evaluated using Hausdorff distance *(H)* between GMM clusters. Briefly, we first calculate all pairwise distances between GMM clusters using Bhattacharyya distance (*B*) for multivariate normal distributions as

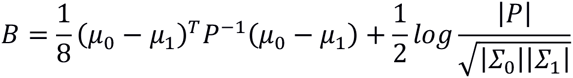

where

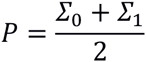

then, as GMM are not ordered, we take the Hausdorff distance *(H)* as the maximum between the row-wise and column-wise minimum of the pairwise distances between two GMM sets. ROC analysis on music distances was performed over the value of *D,*defined as

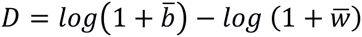

where

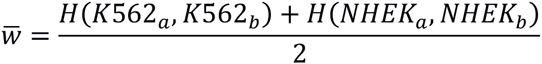

is the average of distances between replicates and

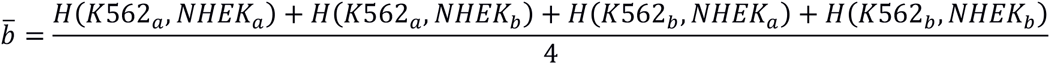

is the average of pairwise distances among different cell lines.

### Assessment of differentially expressed genes

RNA-seq tags were aligned to reference genome using STAR aligner (25). Read counts over RefSeq intervals were extracted using bedtools. Discrete counts were normalized with TMM (26), differential gene expression was evaluated using the voom function implemented in limma (27) with a simple contrast between two cell lines. Genes were considered differentially expressed under a *p*-value lower than 0.01 and absolute logarithm Fold Change higher than 1.

### Cluster analysis

Cluster analysis was performed on replicate 1 of K562 dataset. We calculated all pairwise Hausdorff distances among genomic loci as defined above. Data were clustered using the Ward method. Enrichment analysis was performed using Enrichr (28). Word clouds were created with world_cloud python package (https://github.com/amueller/wordcloud) using text description of ontologies having positive Enrichr combined score. Differential expression among clusters was evaluated using Mann-Whitney U-test.

## Acknowledgements

Authors would like to thank all collaborators and relatives who kindly sacrificed their time to listen to music generated while this work was developed.

